# Cystic fibrosis transmembrane conductance regulator (CFTR) inhibition results in mucus accumulation in human airway epithelia Calu-3 cells: Experimental and Machine Learning Studies

**DOI:** 10.1101/2020.05.26.117853

**Authors:** Brandon S. Laethem, Kenneth T. Lewis, Rafael Ramos, Xia Hou, Fei Sun, Douglas J. Taatjes, Bhanu P. Jena, Suzan Arslanturk

**Author notes:** Correspondence (B.P.J); (S.A). These authors contributed equally to the study.

## Abstract

Porosomes are cup-shaped lipoprotein structures at the cell plasma membrane involved in fractional release of intra-vesicular contents during secretion. At the base of the porosome facing the cell cytoplasm, secretory vesicles dock, fuse and swell, to release intra-vesicular content during secretion. Earlier studies demonstrate the cystic fibrosis trans-membrane conductance regulator (CFTR) associated with the porosome in human airways epithelial Calu-3 mucous-secreting cells, suggesting its possible involvement in porosome-mediated mucus secretion. The current study was undertaken to test this hypothesis. Electron microscopy followed by morphometric analysis using manual and computational machine learning approaches were used to assess changes in secretory vesicle size and content, following stimulation of secretion in the absence and presence of CFTR inhibitors. Results from the study demonstrate that stimulated Calu-3 cells pre-exposed to CFTR inhibitors, demonstrate attenuation of secretory vesicle swelling and the release of mucin. Consequently, accumulation of intracellular mucin is observed in cells exposed to CFTR inhibitors. These results further suggest that mucin secretion from Calu-3 cells involve CFTR both at the secretory vesicle membrane to regulate vesicle volume and hydration, and at the porosome to facilitate mucin hydration and secretion. These new findings progress our understanding of the involvement of CFTR on mucus hydration and secretion, providing critical insights into the etiology of CF disease.

## 1. INTRODUCTION

All living organisms, from yeast to cells in humans depend on secretion to release intracellular products beyond the confines of the cells outer membrane. In mammalian cells, secretion involves the docking, swelling and transient fusion of secretory vesicles at cup-shaped plasma membrane lipoprotein structures called porosomes, for the regulated expulsion of intra-vesicular contents to the interstitium [1-28]. Secretory vesicles dock and fuse [29-37], and undergo swelling involving the transport of water and ions into the secretory vesicle [38-44]. Vesicle swelling result in hydration and a buildup of intra-vesicular pressure, required for content expulsion during cell secretion. Greater pressure translates to increased intra-vesicular content release. This mechanism of secretion retains the integrity of the secretory vesicle membrane and the cell plasma membrane, while precisely regulating intra-vesicular content release [45-47] during cell secretion. The cup-shaped porosomes range in size from 15 nm in neurons and astrocytes, to 100-180 nm in endocrine and exocrine cells. Porosomes in Calu-3 cells measure approximately 100 nm and are composed of nearly 34 proteins among them the cystic fibrosis trans-membrane conductance regulator (CFTR) [48], compared to the nuclear pore complex of similar dimension and nearly 1,000 protein molecules [49]. Human airways epithelium is coated with a thin film of mucus, composed primarily of mucin. Mucus is continually moved via ciliary action and cleared. In addition to lubricating the airways, mucus serves to trap foreign particles and pathogens, assisting in keeping the airways free of infection. Mucus hydration has been recognized as a problem in cystic fibrosis (CF), and therefore we posit that altered mucin secretion either due to increased viscosity of the secreted mucin or attenuated secretion, would results in the inability of the cilia to propel mucin, leading to its stagnation in the airways and to infections observed in CF disease. To test this hypothesis, the human lung epithelia cell line Calu-3 was used. Calu-3 cells were stimulated in the presence and absence of the CFTR inhibitors 172 and GlyH101 [50], to determine changes in secretory vesicle size and content (filled vs empty). Transmission electron microscopy followed by morphometry of the electron micrographs performed both manually and using computation machine learning approach, were used to determine changes in mucin secretion.

## 2. MATERIALS AND METHODS

### 2.1 Calu-3 cell culture

Airways epithelial Calu-3 cells are derived from a human lung adenocarcinoma [51]. The cells were grown in Dulbecco’s Modified Eagle Medium: Nutrient Mixture F-12 (DMEM/F-12) (Invitrogen) containing 15% fetal bovine serum. Cells were incubated in a humidified atmosphere at 37°C and 5% CO_2_.

### 2.2 Stimulation of Calu-3 cells

Stimulation experiments were performed on Calu-3 cells grown to confluency in an air-liquid interface on permeable filter supports. Calu-3 cells were seeded on Costar Transwell permeable supports (Costar, Corning) in DMEM:F12 (Sigma-Aldrich, St.Louis.MO, USA), supplemented with 10% Fetal Bovine Serum, 1% Pen/Strep, 10μg/ml Gentamycin. for establishing the air-liquid interface cultures. For Ussing chamber experiments, cells were cultured at 37° C and 5 % CO2 in Ringer’s buffer (120mM NaCl, 25mM NaHCO_3_, 1.2mM CaCl_2_, 1.2mM MgCl_2_, 1.2 KH_2_PO_4_, 0.8mM K_2_HPO_4_ and 11.2 glucose) that was supplemented with 15 % fetal calf serum (FCS), 500 U/ml penicillin and 50 μg/ml. The medium was added to both the apical and basolateral compartments. An air-liquid interface culture was initiated by removing medium from the apical compartment on the second day after plating cells and the apical compartment containing cells was exposed to air. The medium (0.5ml) in the basolateral compartment was changed every 2-3 days. After approximately 11 days of culturing at air-liquid interface, tight conjunctions between cells are established. The filter with cells is then transferred and mounted in the Ussing chamber holder. Ringer’s buffer is added to the apical side of the chamber and low Ringer’s (60mM Sodium Gluconate, 25mM NaHCO_3_, 1.2mM CaCl_2_, 1.2mM MgCl_2_, 1.2 mM KH_2_PO_4_, 0.8mM K_2_HPO_4_ and 11.2 mM glucose) to the basolateral compartment. *I*_sc_ was measured by voltage clamping. Once a stable basal point was established, 10μM Forskolin was added to the basolateral compartment to induce channel opening. When the CFTR channels reach a stable activation threshold, 10μM inhibitor GyH101 was added to inhibit channel activity.

### 2.3 Electron microscopy

Transmission electron microscopy of forskolin-stimulated Calu-3 cells in the presence and absence of the CFTR inhibitors GlyH101 and Inh172, were prefixed for 2h in 4% PFA and then stored in 2% PFA until further processing was performed. Briefly, cells were further fixed in half-strength Karnovsky’s fixative (1.5% glutaraldehyde, 1.0 % formaldehyde in 0.1M cacodylate buffer, for 80 minutes at 4oC, followed by post-fixation for 1h at 4°C in 1% OsO_4_ in 0.1 M cacodylate buffer. The sample was then dehydrated in a graded series of ethanol through propylene oxide and infiltrated and embedded in Spurr’s resin. Ultrathin sections were cut with a diamond knife, retrieved onto grids, and contrasted with alcoholic uranyl acetate and lead citrate. Grids were viewed with a JEOL 1400 transmission electron microscope (JEOL USA, Inc., Peabody, MA) operating at 80 kV, and digital images were acquired with an AMT-XR611 11 megapixel ccd camera (Advanced Microscopy Techniques, Danvers, MA).

### 2.4 Manual and deep learning algorithms for morphometric analysis

*ImageJ* image processing software was used to manually demarcate the outline of cells and secretory vesicles within electron micrographs of forskolin-stimulated Calu-3 cells in the presence and absence of the CFTR inhibitors GlyH101 and Inh172, and under identical magnification, were used to determine changes in vesicle size and contents within cells. This manual morphometric measurement was complemented by a machine-learning framework developed to understand the morphometry and content of the secretory vesicles in the presence of the two CFTR inhibitors (GlyH101 and Inh 172). In particular, a deep learning segmentation method based on a ResNet-101 network architecture [52-54] was employed, that allows the automatic detection of secretory vesicles.

## 3. RESULTS AND DISCUSSION

Ussing chamber studies demonstrate the presence of a functional CFTR in Calu-3 cells [Figure 1]. When exposed to 10μM Forskolin, Calu-3 cells release chloride, which is attenuated in the presence of the two CFTR inhibitors (GlyH101 and Inh 172). Manual demarcation of cells and secretory vesicles in electron micrographs of Calu-3 cells under identical magnification demonstrate a loss of intra-vesicular contents following stimulation of secretion. In contrast, cells pre-exposed to CFTR inhibitors reflect little loss of vesicle contents following stimulation of secretion [Figures 2-4].

**Figure 1.**
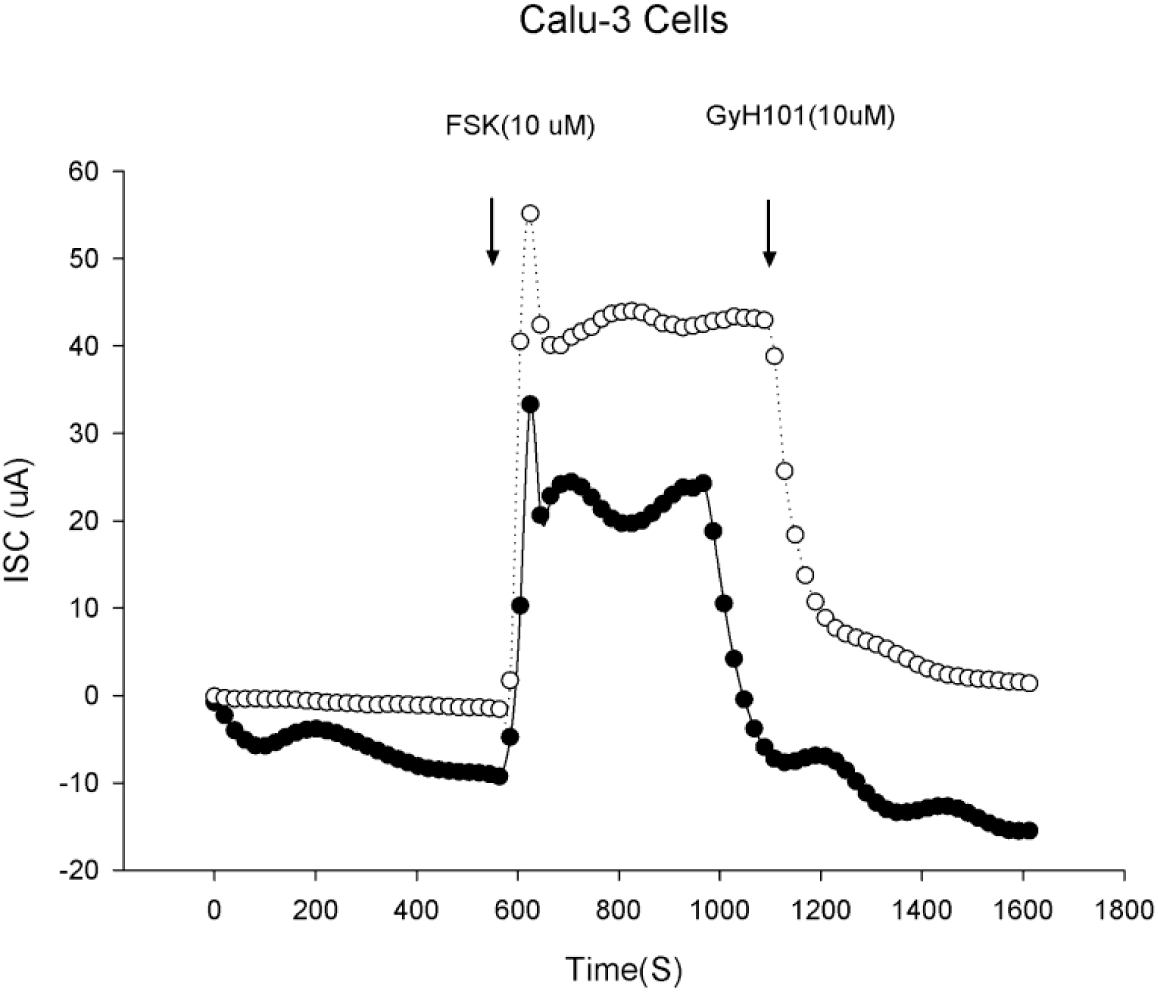
Ussing chamber experiments showing Forskolin-stimulated chloride release from Calu-3 cells which is inhibited in the presence of the CFTR inhibitor GlyH-101. Note that the two separate experiments demonstrate similar stimulation and inhibition profiles.

**Figure 2.**
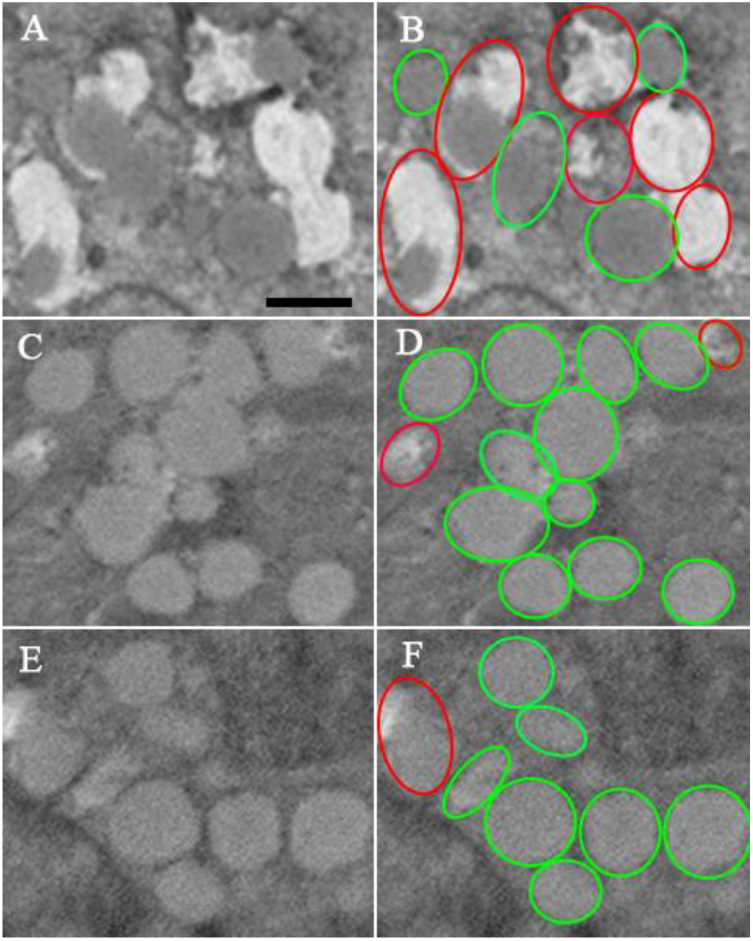
CFTR inhibitors 172 and GlyH-101 inhibit forskolin-stimulated secretion of intra-vesicular mucin from Calu-3 cells. A and B are from the control cell, C and D are from cells exposed to 172, and E and F are from cells exposed to GlyH-101. The vesicles outlined in red are the partial/ empty vesicles and those outlined in green are the filled vesicles. *Scale bar = 500 nm*.

**Figure 3.**
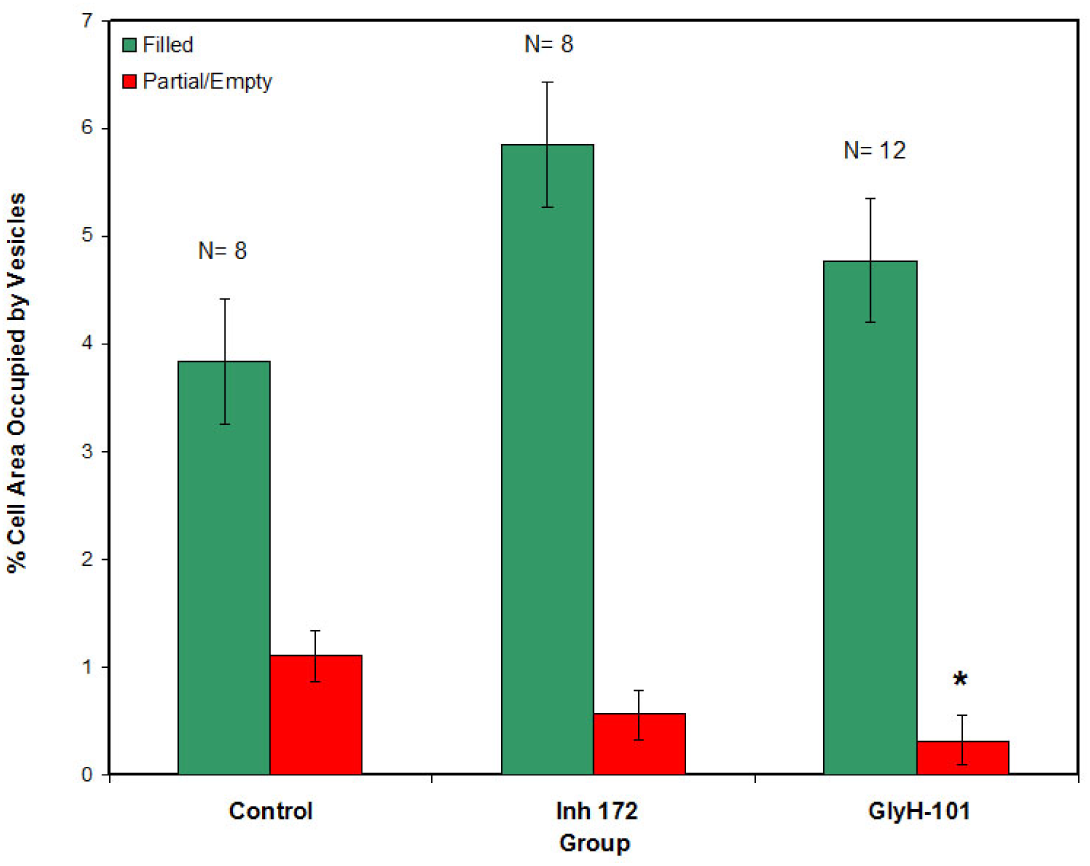
Manual morphometry demonstrates that exposure of Calu-3 cells to the CFTR inhibitors GlyH-101 inhibit stimulated secretion from Calu-3 cells. Note the significant (*p<0.05) loss of secretion and retention of intra-vesicular contents in Calu-3 cells exposed to GlyH-101.

**Figure 4.**
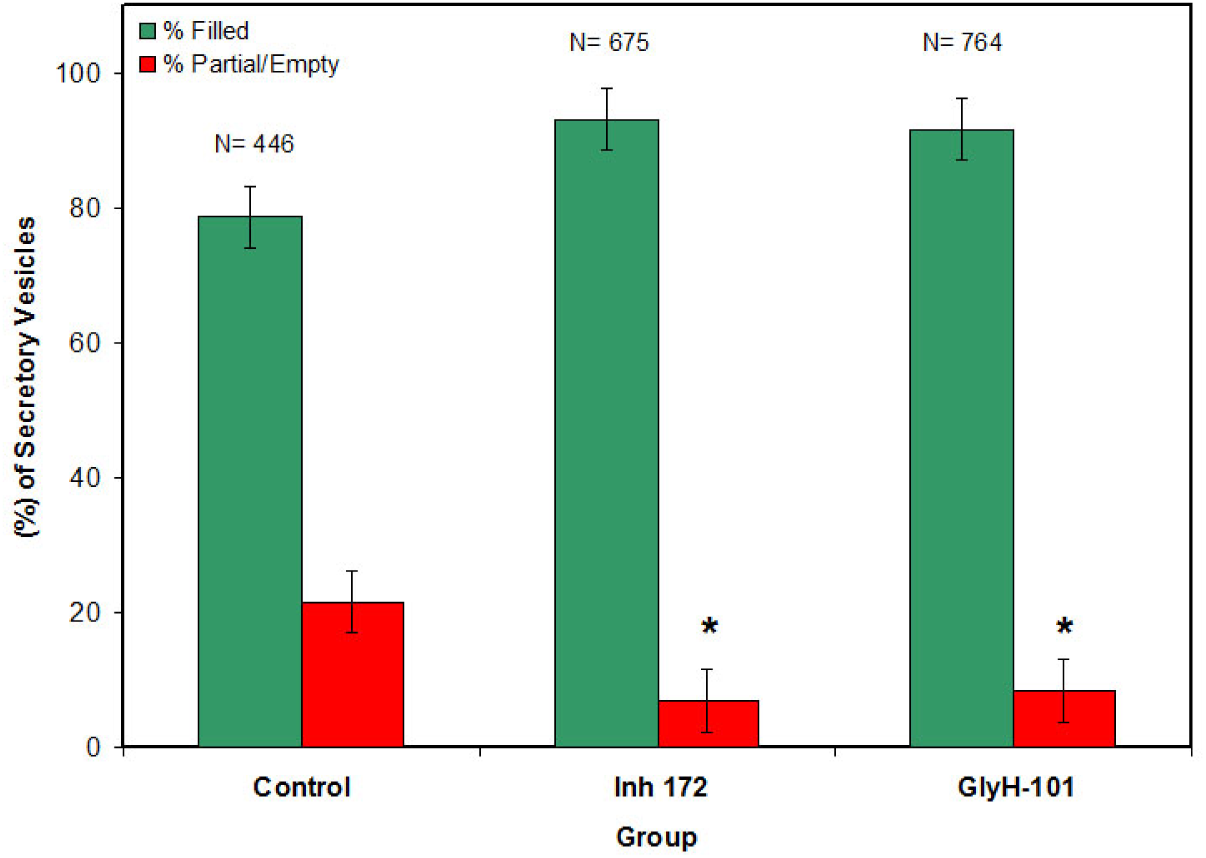
Manual morphometry of secretory vesicles demonstrates that exposure of Calu-3 cells to the CFTR inhibitors Inh172 and GlyH-101, inhibits stimulated secretion from Calu-3 cells. Note the significant (*p<0.05) drop in the presence of partially empty vesicles in Calu-3 cells exposed to both the CFTR inhibitors.

Here, we built a framework [Figure 5] to understand the morphometry and content of the secretory vesicles in the presence of the two CFTR inhibitors (GlyH101 and Inh 172). In particular, we have built a deep learning based segmentation method on a ResNet-101 network architecture [52-54] that allows the detection of the objects of interest, i.e. the secretory vesicles. Object segmentation refers to the task of grouping pixels that define an object and requires the annotation of several foreground and background objects to perform a supervised learning task. The partial and filled secretary vesicles in multiple Calu-3 cells are annotated manually and pixel-by-pixel as foreground objects. During training, the model “learns” to distinguish the foreground objects from the background. A separate model is trained for control and each CFTR inhibited cells. Once the fully trained ResNet-101 models are constructed based on the labeled data, new Calu-3 cell images are fed into the network to automatically annotate each pixel and determine the group of pixels representing the vesicles. Figure 6 shows the segmentation of partial and filled secretory vesicles on control and CFTR inhibited images using our computational approach. Next, the features associated with each secretory vesicle are estimated using computational image analysis. The extracted features are; the average amount of gray intensity, minimum intensity, maximum intensity, standard deviation of the intensity, number of pixels within the object area, 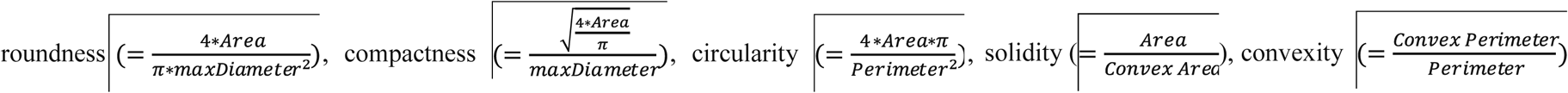 the x- and y-coordinates of the center of each object.

**Figure 5.**
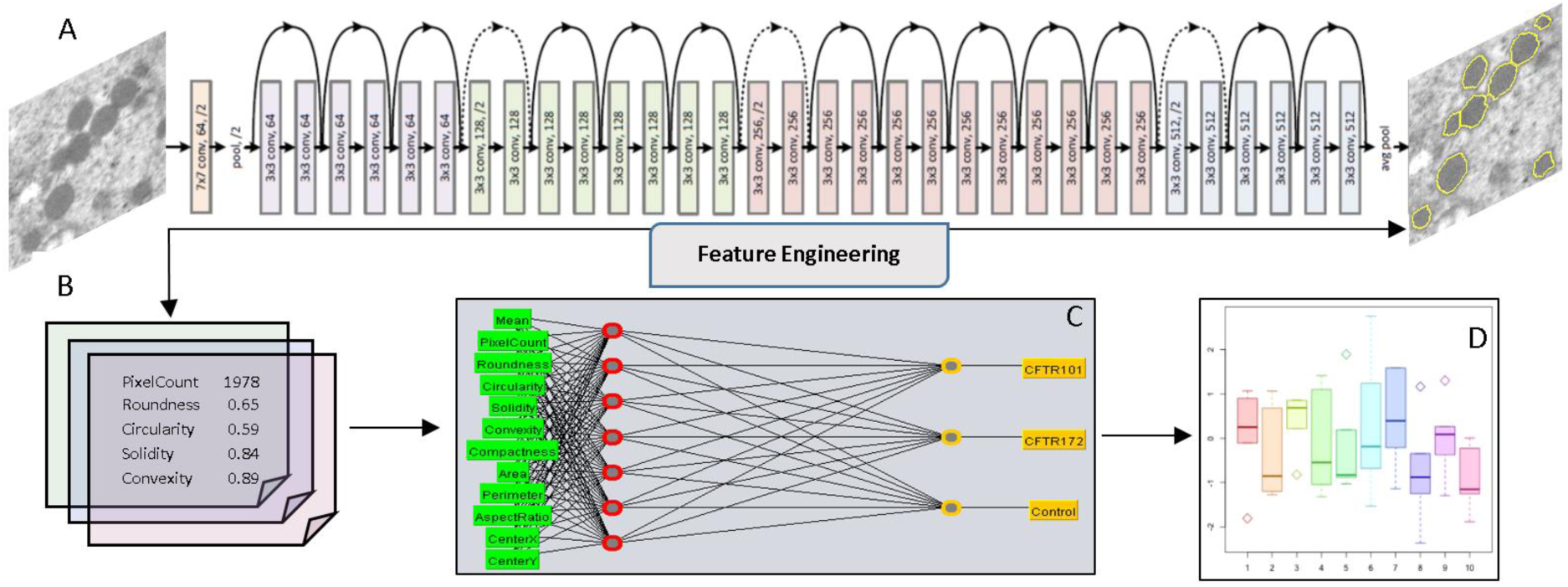
Computational Analysis Framework. **A**. Deep Learning based segmentation usi ng a ResNet-101 network architecture; **B**. Parameters obtained through computational im age analysis and feature engineering; **C**. The artificial Neural Network (ANN) architectur e that differentiates vesicles in the control and CFTR inhibited Calu-3 cells using paramet ers obtained in Step B; **D**. Identification of significant parameters associated with the ch anges in secretory vesicle content in control, and in the presence of the two CFTR inhib itors.

**Figure 6.**
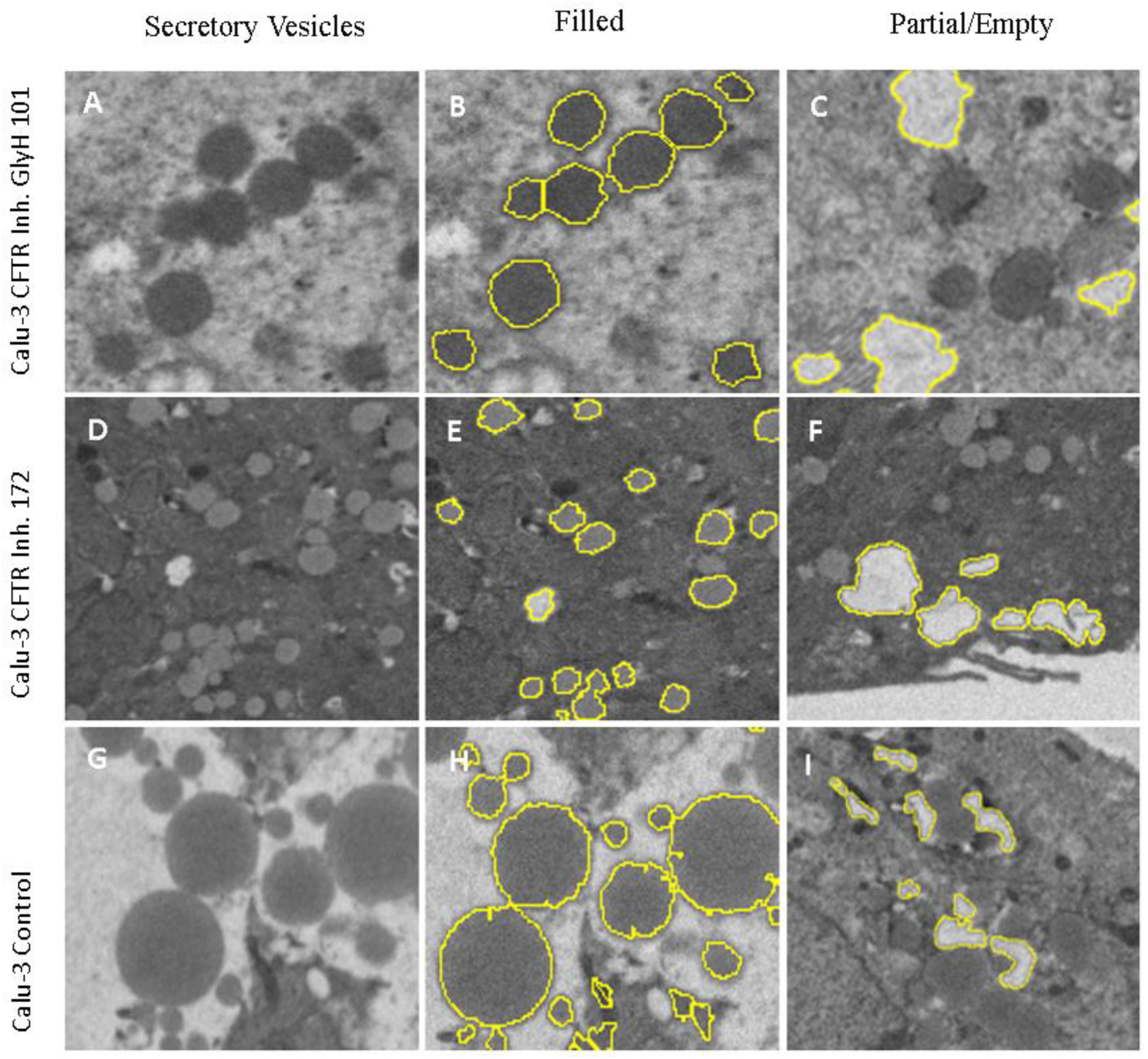
Filled and partially filled/empty secretory vesicles (scale bar = 2 um). **A**. Gly H-101 cell with filled and partially filled/empty vesicles; **B**. GlyH-101 filled vesicle seg mentation; **C**. GlyH-101 partial vesicle segmentation; **D**. Inhibitor 172 cell with filled an d partially filled/empty vesicles; **E**. Inhibitor 172 filled vesicle segmentation; **F**. Inhibitor 172 partially filled/empty vesicle segmentation; **G**. Calu-3 control (unexposed to CFTR inhibitors) cell with filled and partially filled/empty vesicles; **H**. Filled vesicle segmentation in control; **I**. Partially filled/empty vesicle segmentation in control.

The extracted features are then fed into an artificial neural network (ANN) that differentiates the vesicles in the control and CFTR inhibited Calu-3 cells. The architecture of the ANN is illustrated in Figure 5c. Once a fully trained ANN is constructed, the backpropagation (i.e. differentiation of output classes with respect to input features) of the ANN would enable the assignment of contribution scores to each input feature. Here, the prediction score is distributed to each input feature in proportion to its contribution to the prediction with respect to a certain baseline class (i.e. control, GlyH101 and Inh 172). Those inputs with high prediction scores are considered to be significant markers as they contribute decisively on the selected class.

The ANN model is tested on a 10-fold cross-validation set and the model performance is assessed through the metrics defined in Table 1. The accuracy of the validation set is estimated to be 93%.

**Table 1.** Detailed Accuracy of the ANN by Class

|  | **TP Rate** | **FP Rate** | **Precision** | **Recall** | **F-Measure** | **ROC Area** | **Class** |
| --- | --- | --- | --- | --- | --- | --- | --- |
|  | 0.949 | 0.081 | 0.943 | 0.949 | 0.946 | 0.975 | **CFTR101** |
|  | 0.91 | 0.011 | 0.924 | 0.91 | 0.917 | 0.992 | **CFTR172** |
|  | 0.91 | 0.034 | 0.916 | 0.91 | 0.913 | 0.972 | **Control** |
| **Weighted Avg.** | 0.933 | 0.059 | 0.933 | 0.933 | 0.933 | 0.976 |  |

The significant features extracted through backpropagation are the average amount of grey intensity area, circularity and x-coordinates of the center of each vesicle (denoted as CenterX). Figure 7A-D shows the significant feature distributions of the vesicles by class.

Our computational results show that circularity and CenterX can further improve the classification accuracy by distinguishing between GlyH101 and Inh 172 cells (Figure 7b-c). In agreement with data obtained using manual morphometric analysis, our computational measurements demonstrate a reduction in the surface area of secretory vesicles in CFTR inhibited Calu-3 cells. This may be due in part to the prevention of ion and water entry into vesicles for hydration and consequent vesicle swelling required during cell secretion [38-44]. These results further suggest that mucin secretion from Calu-3 cells involve CFTR function, both at the secretory vesicle membrane to regulate vesicle water entry and hydration, and the additional hydration of mucin at the porosome complex to facilitate secretion.

**Figure 7.**
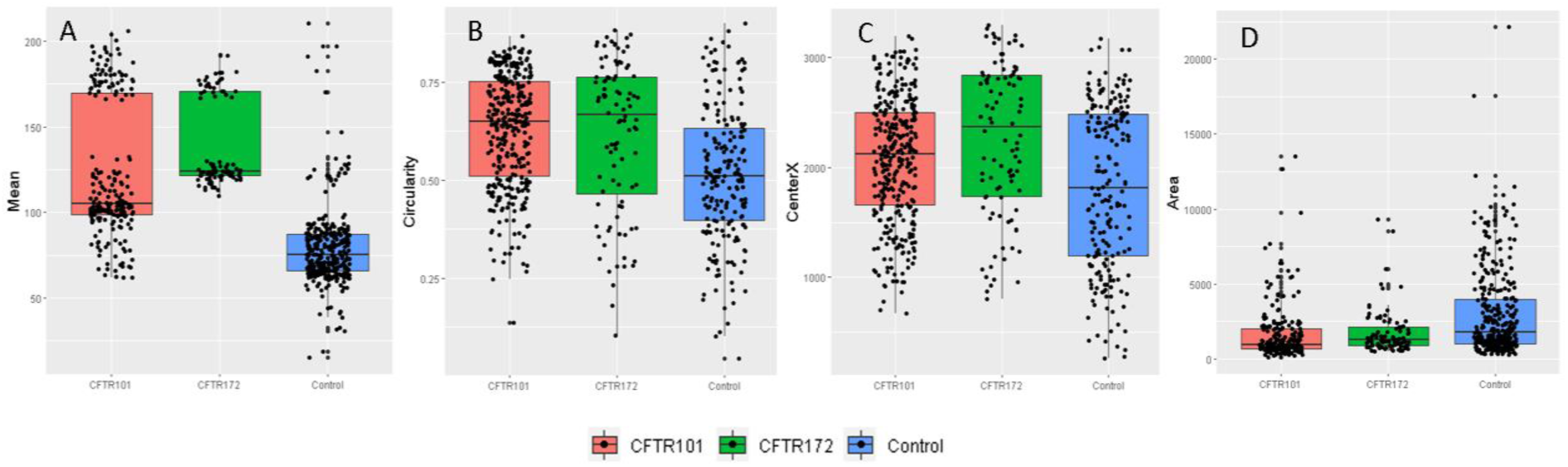
Filled and partially filled/empty secretory vesicle grey intensity (or electron dense i.e., content), circularity, X-coordinates and area, in control Calu-3 cells and cells exposed to CFTR inhibitors GlyH-101 and Inh172. **A**. The average amount of gray intensity **B**. Circularity, **C**. X-coordinates of the center of each vesicle (Center X) and **D**. Area distributions of the vesicles by class. The computational analysis have demonstrated that changes in the aforementioned significant features can successfully lead to the differentiation of control and CFTR-inhibited cells. In particular, due to the higher percentage of empty and partially empty secretory vesicles in control cells (also demonstrated through the manual morphometry analysis in Figure 4), the average amount of gray intensity in control cells is observed to be less than the CFTR inhibited cells (Figure 7A).

## Author Contributions

B.P.J. developed the idea. B.P.J. and S.A. wrote the paper. B.S.L. and K.T.L. performed the experiments. B.S.L., K.T.L. and R.R. carried out manual morphometric analysis and prepared some of the figures. D.J.T performed the electron microscopy. X.H and F.S performed the Ussing chamber assays. S.A performed machine learning analysis on the EM data. All authors participated in critical reading and discussion of the manuscript.

## Acknowledgements

Work presented in this article was supported in part by the National Science Foundation grants EB00303, CBET1066661 (BPJ).

